# *SOX2* dosage sustains tumor-promoting inflammation to drive disease aggressiveness by modulating the *FOSL2/IL6* axis

**DOI:** 10.1101/2022.06.09.495487

**Authors:** Abdel Jelil Njouendou, Arnol Auvaker Zebaza Tiofack, Rovaldo Nguims Kenfack, Sidonie Noa Ananga, Esther H. M. Dina Bell, Gustave Simo, Jörg D. Hoheisel, Jens T. Siveke, Smiths S. Lueong

**Author notes:** Corresponding author, Dr. Smiths S. Lueong.

## Abstract

**Background:** Inflammation is undoubtedly a hallmark of cancer development. Its maintenance within tumors and subsequent consequences on disease aggressiveness is less understood.

**Methods:** Multi-omic analyses of 27 (~ 5000 samples) entities from the TCGA, GEO and in-house data was performed to investigate the molecular determinant of tumor aggressiveness. Using molecular loss of function data, the mechanistic underpinnings of inflammation-induced tumor aggressiveness was addressed.

**Results:** The data revealed a significant association between somatic copy number alterations (sCNA) and tumor aggressiveness, with amplification of the transcription factor *SOX2* being the most important feature among novel and known aggressiveness-associated genes such as *ZIC5* and *MYEOV.*

Mechanistically, *SOX2* regulates a group of aggressiveness-related genes including the *AP1* transcription factor *FOSL2* to sustain pro-inflammatory pathway such as *IL6-JAK-STAT3, TNFA* and *IL17* signaling pathways. Prolonged inflammation induces immunosuppression and further leads to activation of cytidine deamination and consequential DNA damage evidenced by enrichment in cytidine deamination mutational signatures in aggressive tumors.

The resulting DNA damage affects tumor suppressor genes such as *TP53,* which was the most mutated gene in aggressive tumors compared with less aggressive tumors (38% vs 14%), thereby releasing cell cycle control. This was exemplified in Glioblastoma multiform, where *TP53* and *IDH1* mutations are predominant. *IDH1* mutations were almost only seen in younger patients (>45 years, > 90%) and may explain the previously reported favorable prognosis.

**Conclusion:** Taken together, our data demonstrate the implication of *SOX2* in promoting DNA damage and genome instability by sustaining inflammation via *FOSL2/IL6,* resulting in tumor aggressiveness.

## Introduction

Genomic instability is undoubtedly a major hallmark of cancer development, as it drives tumor heterogeneity and supports the adaptation of cancer cells to stress conditions leading to the malignant behavior and therapy resistance (1, 2). Oncogenic driver mutations or inactivation of DNA repair genes (3) are amongst the early events in malignant transformation and are involved in the process of tumorigenesis (4). Other alterations such as deletions, amplifications, fusions and translocations are not of dismal consequences. The extent to which different genomic alterations shape the final tumor cell phenotype is context-dependent. Precision oncology seeks to tailor patient treatment schemes to specific molecular portraits (5). Genomic instability therefore represent a considerable challenge to personalized oncology.

Following malignant transformation, tumors exhibit diverse clinical phenotypes even within the same entity. Some cancers such as pancreatic ductal adenocarcinoma (PDAC), glioblastoma multiform (GBM) and ovarian cancers (OV) are notoriously aggressive and lethal (6–8). In PDAC and GBM, the mesenchymal phenotypes are even more aggressive (9, 10) and similar observations are reported in triple negative breast cancer (TNBC) (11). These aggressive tumors are generally undifferentiated and plastic (9, 12). Several molecular drivers have been proposed to drive tumor aggressiveness in different cancer entities, such as *FOSL1* expression in brain cancer (9), stromal stiffness, miRNA silencing (7) as well as *MYC* expression and *KRAS* gene dosage in pancreatic cancer (13–15). In breast cancer, high mutation rates have been associated with aggressive tumor phenotypes (16). Gene dosage can therefore modulate disease phenotype or contribute in shaping it. In prostate cancer for example, somatic copy number alterations (sCNA) in the *BNIP3L, WWOX,* and *GATM*genes are responsible for tumor aggressiveness (17). In effect, alterations in gene dosage might alter expression and thus cellular homeostasis, leading to tumor development or tumor progression and therapy resistance.

Apart from sCNA, translocations, fusions and extra chromosomal gene amplifications equally play substantial roles in defining the net clinical disease phenotypes (18–22). Furthermore, early onset of cancer, at least in some cancer entities such as colorectal and gastric cancers is associated with aggressive disease profiles (23, 24). Lastly, inflammatory tumor microenvironments (TME) promote cancer metastasis and aggressiveness (25–27). In effect, inflammation activates cytidine deamination via proinflammatory transcription factors such as NFkB leading to DNA double strand breaks and tumorigenic pathway activation by supporting the susceptibility to mutagenesis in several epithelial organs (28)

A comprehensive analysis of how these different molecular factors synergize to influence disease phenotype is still missing, although it can tremendously improve the management of aggressive tumors. In the present study, an integrative multi-omic pancancer analysis of data from the cancer genome atlas (TCGA) and the gene expression omnibus (GEO) as well as in-house generated data was undertaken to identify the major genomic determinants of cancer aggressiveness and their mechanistic underpinnings. The data analysis uncover an association between somatic copy number amplifications and disease aggressiveness. We demonstrate that *SOX2* copy number amplification and corresponding transcriptional upregulation is the most significant alteration and investigate its mechanism. Mechanistically, *SOX2* regulates the API transcription factor *FOSL2,* which in turns support an inflammatory TME by promoting *TNFA, IL6-JAK-STAT3* and other proinflammatory signaling pathways to induce immunosuppression and stimulate genomic instability by activation of cytidine deamination.

## Methods

### Data mining

Molecular and clinical data from 27 cancer entities from the TCGA as analyzed. A list of all cancer entities included is presented in **supplementary table 7**. For all 27 cancer entities included in this study, molecular and clinical data was downloaded from the cancer genome atlas (TCGA). The TCGAbiolinks Bioconductor package was used for data download. sCNA data was Affymetrix SNP 6.0 array data from the legacy database (hg19 genome assembly), downloaded using the GDCquery, download and prepare commands of the TCGAbiolinks package. Copy number segmentation data was downloaded only for primary tumors, by specifying the sample.type to “Primary Tumors” while data category and datatype were respectively set to “copy number variation and “nocnv_hg19.seg”, respectively. When the total number of samples for a given cohort was more than 250 cases, only the first 250 samples were downloaded by sub setting the output from the GDCquery command.

For gene expression data, the legacy database was equally used to download illumina HiSeq gene expression quantification data from primary tumors. For each entity, RNA-seq data from all available samples was downloaded.

MAF files containing somatic mutation data from the harmonized data base was used. Somatic mutation data used was preprocessed using the TCGA “muse” pipeline. MAFtools was used to estimate tumor mutational burden as well as mutational signatures

Gene expression and ChIP-seq data derived from *SOX2* knockdown cells was downloaded from the gene expression omnibus (GEO). The downloaded data are available through the following accession numbers: E-GEOD-46837, GSE166184, GSE166183. Additional *SOX2* and *FOSL2* ChIP-seq data was downloaded from the encyclopedia of DNA elements (ENCODE) derived from the breast cancer cell line MCF7. Gene expression data from pancreatic normal and disease tissue was generated in-house and is available under the accession number E-MTAB-1791.

### sCNA data analysis

Following data download, sCNA data was analyzed using the gaia Bioconductor package. To this end, probe meta files for the hg19 genome assembly was downloaded from the Broad institute and sex chromosome names were converted to the respective numbers (X = 23 and Y = 24). After duplicate removal, the marker matrix was filtered for common copy number alterations (CNVs) that are usually present in normal samples. The filtered marker matrix was then used to create a marker object. Copy number variation data from each entity was then downloaded as described above and converted into a matrix. A copy number segment threshold of −0.3 and 0.3 was used to delineate copy number deletions (sCNAdel) and somatic copy number amplifications (sCNAmp), respectively. Sex chromosome names were equally replaced with respective numbers to match the marker object. Somatic copy number alterations were then identified by running the gaia command on the marker object, the processed CNA matrix and for the number of samples in the CNA matrix. sCNA data was annotated using the GenomicRanges and biomaRt Bioconductor packages. The same packages were equally used to intersect and annotate sCNA from different cancer entities. The circlize package was used to generate circus plots for sCNA and somatic mutation data

### Analysis of gene expression and ChIP-seq data

Raw gene expression data from the TCGA was normalized and quantile filtered using the EDASeq Bioconductor package. The filtered data was then converted into a DGEList object for downstream analysis using the edgeR package. Differentially expressed genes were selected by defining a log2 fold change of 0.9 in the TCGA data and 1 for cell line-derived data. For all data sets, the false discovery threshold was set to < 0.05. The list of all differentially expressed genes was then used to subset the gene expression matrix from generation of heatmaps. The gene expression deconvolution tool, TIMER (http://timer.cistrome.org/) was used to determine the proportions of immune cell infiltration in gene expression data.

For ChIP-seq data anlysis, fastq files were assessed for their quality using the FastQc tool and trimmed with trimmomatic. The trimmed data was the mapped to the hg38 human genome assembly using bwa short read mapper. The mapped reads were filtered and used for peak calling with MACS. Bedgraph and narrow peak files were exported and annotated using the ChIPpeakAnno Bioconductor package. ChIP-seq peaks were displayed using the bioconductor package trackViewer.

Somatic mutations and mutational signatures were analyzed using Bioconductor package MAFtools. The gene TTN was excluded from all oncoprints, meanwhile only the top5 mutational signatures were retrieved

Gene set enrichment analysis was performed using the GSEA algorithm from the Broad institute for the hallmarks gene set, while the Bioconductor package PathfindR was used for KEGG pathway analysis. The significance threshold for GSEA and KEGG analysis was set to < 0.05 (29). Genes associated with aggressive cancer phenotype were determined using the boruta feature selection package.

### Statistical analysis

Patient samples were classified as early onset, if the patient was diagnosed before or at the age of 45 years (mean age at diagnosis for all entities – 1 standard deviation). They were otherwise considered as late onset. Aggressive cancers (poor outcome) were considered to be those with more than 40% of cases declared dead at the end of each study, meanwhile cancers with less than 20% of cases declared dead were considered as less aggressive (better outcome). All patients without age or gender information were excluded from the analysis. Similarly, for survival analysis, all patients with missing data for time to last follow-up or vital status were equally excluded. Survival analysis was performed using the survival and survminer R packages. Cut-off threshold for dichotomization of gene expression data was determined using the surv_cutpoint function of the survminer package. All analyses were performed with the R environment or using Graphpad prism version 8.0.0 for Windows, (GraphPad Software, San Diego, California USA, www.graphpad.com”).

## Results

### Somatic copy number alterations are associated with tumor aggressiveness

We investigated the molecular traits of tumor aggressiveness in 27 cancer entities from the TCGA. Molecular data from highly aggressive cancers (>40% fatalities) and less aggressive cancers (< 20% fatalities) were compared. Based on this criteria, 10 cancers including: GBM, PDAC, OV, SKCM, CHOL, BLCA, LUSC, HNSC, ESCA and STAD were categorized as highly aggressive, meanwhile PRAD, THCA, THYM, TGCT, KICH KIRP and BRCA were categorized as less aggressive (**figure 1a**). More than 50% of patients with aggressive tumors had died within 2500 days of follow-up meanwhile more than 90% of patients with less aggressive tumors were alive after 10000 days of follow-up (**figure 1b**). Mean age at diagnosis of all cancer entities was around ~ 58 ± 13 years (**figure 1c**). We defined early onset of cancer as cancer diagnosed before 45 years of age (mean – 1SD). A cancer was considered to have substantial early onset, if more than 10% of cases were diagnosed before 45 years of age. There was no direct relationship between age at diagnosis an percentage fatality, as some less aggressive tumors such as PRAD was exclusively in older patients (**figure 1d**). Correlation analyses revealed that disease fatality was positively correlated with the total number of sCNA but not tumor mutational burden (TMB) (**figure 1e**). In some highly aggressive tumors such as SKCM, high tumor mutational burden was observed both in young and older patients at comparable proportions. However, comparable but lower TMB was observed in GBM, suggesting other possible causes of aggressiveness (**figure 1f**). Generally, copy number deletions were more prominent than amplifications (**figure 1g & 1h**) and there were more somatic copy number alterations in highly aggressive than in less aggressive tumors (**supplementary figure 1a, 1b, & 1c (left panel)**). No such obvious trend was observed for TMB, except in SKCM (**supplementary figure 1 right panel**).

**Figure 1:**
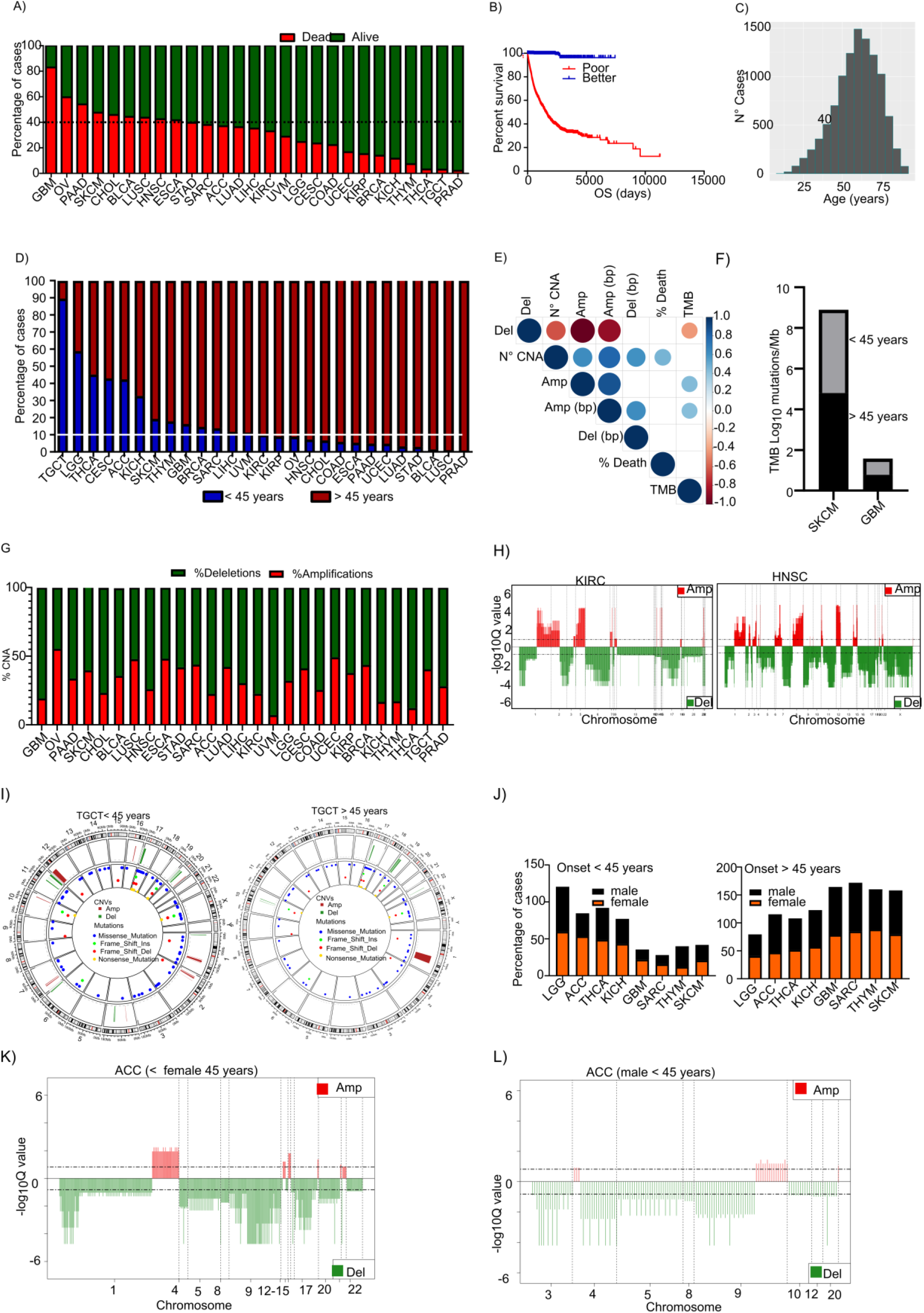
Somatic copy number alterations are associated with cancer aggressiveness. **A**) A bar plot presenting the percentage of cancer-related casualties in 27 cancer entities from the TCGA. Each bar represent a cancer entity and the green color represent the percentage of all patients who were reported alive at the last follow-up, while the red bars represent the percentage of patients who were confirmed dead at the last follow-up. **B**) A Kaplan-Meier overall survival curve for patients from the first 10 poor outcome cancer entities (red) and the last 5 better outcome entities (blue) (from figure 1a). The survival time represent the time from first diagnosis to last follow-up. **C**) A histogram showing age distribution across all analyzed 27 cancer entities from the TCGA. The average age at first diagnosis for all cancer entities analyzed was ~58 years with a standard deviation of ~13 years **D**) A bar plot presenting the fraction of patients who were diagnosed with cancer before or after 45 years. The cut-off of 45 years represents the mean age at diagnosis minus one standard deviation around the mean. **E**) A correlation plot showing the correlation between the percentage of dead cases per entity and copy number alterations (CNA) as well as tumor mutational burden (TMB). The number of identified CNA (N° CNA) as well as the number of deletions (Del) or amplifications (Amp) and the number of amplified (Amp bp) or deleted base pairs (Amp bp) are presented. **F**) A bar plot showing the tumor mutational burden in two of the most aggressive entities, where more than 10% of cases were diagnosed before 45 years of age. **G**) A circus plot showing the copy number alterations and mutations in TGCT for patients diagnosed befpre (left panel) and patients diagnosed after (right panel) 45 years of age. TGCT is the entity with the highest number of patients diagnosed before 45 years of age. **H**) A bar plot showing the percentage of amplifications or deletion in each of the 27 cancer entities. **I**) Representative CNA plots showing copy number amplifications or deletions in kidney renal cell carcinoma (left panel) and head and neck squamous cell carcinoma (right panel). **J**) Bar plots showing the percentage of male and female cases diagnosed before the age of 45 years, for all cancer entities with more than 10% of cases diagnosed before 45 years of age. Gender-specific entities are not included here. **K & L**) Gaia can plots for ACC in females and males diagnosed before the age of 45 years, respectively. ACC is one of the entities with higher incidence in females before 45 years of age

### sCNA is associated with cancer onset

We then asked if sCNA as equally associated with cancer onset. To address this, we used data from TGCT with more than 80% early onset. We observed more sCNA in younger patients compared with older patients (**figure 1g**). Given recent reports on gender discrepancies in cancer, we investigated possible gender effects in early cancer onset on other cancers with early onset. In younger patients, the female fraction of ACC and GBM were higher, while more older males had ACC but a comparable distribution of GBM with females. THYM and SKCM were predominant in younger but not older males (**figure 1J**). There was no difference in the TMB for cancers with early onset between males and females (**supplementary figure 1d**). To further establish the involvement of sCNA in cancer onset, we specifically looked at Adrenocortical carcinoma (ACC). We assessed if females diagnosed with ACC before 45 years showed more sCNA alterations than males diagnosed within the same age range. In effect, there were more sCNA in young females than in males (**figure 1k & 1l**). In GBM there was no difference in sCNA in early and late onset of GBM. We further investigate additional layers of omics data in GBM, as it was the most lethal of all cancers.

### *IDH1* and *TP53* mutations are associated with early onset of GBM

To better understand the molecular basis of GBM aggressiveness and onset, we analyzed somatic mutations as well as sCNA and gene expression. Early onset of GBM was associated with high levels of *TP53, IDH1* and *ATRX* mutations (**figure 2a & 2b)**. In late onset cases, *PIK3CA* mutations were observed in 7% of cases, but not in early onset cases. Given that, *IDH1* mutations are often associated with better survival (30, 31); we wondered if this was due its predominance in younger patients. We then compared patient overall survival in *IDH1* mutant and wild type patients with overall survival for early and late onset (**figure 2c & 2d).** Further analysis of sCNA revealed copy number amplifications of specific oncogenic drivers in late onset, which were absent in early onset cases (**figure 2e & 2f**). Precisely, a zoom in on chromosome 3 revealed sCNA amplification around the *PIK3CA* and *SOX2* gene loci in late onset. We then performed differential gene expression analysis between early and late onset of GBM and found only very few differentially expressed genes (**figure 2g**). It is therefore very likely, that early GBM onset is associated with DNA damage while sCNA is more prominent in late onset GBM. In support of this, differences in somatic mutation patterns were equally observed for early and late onset in other tumor entities. For example, in Sarcoma, more than 30% of *TP53* mutations were observed in late onset cases as opposed to less than 10% in early onset (**supplementary figure 2a & 2b**). Similarly, in Thymoma, more than 60% of cases showed mutations in *GTF2I,* while only 18% of early onset cases were affected by this mutation (**supplementary figure 2c & 2d**). Lastly, 30% of cases with late onset of TGCT harbored *KIT* mutations, while only 8% of early onset cases were found with *KIT* mutations.

**Figure 2:**
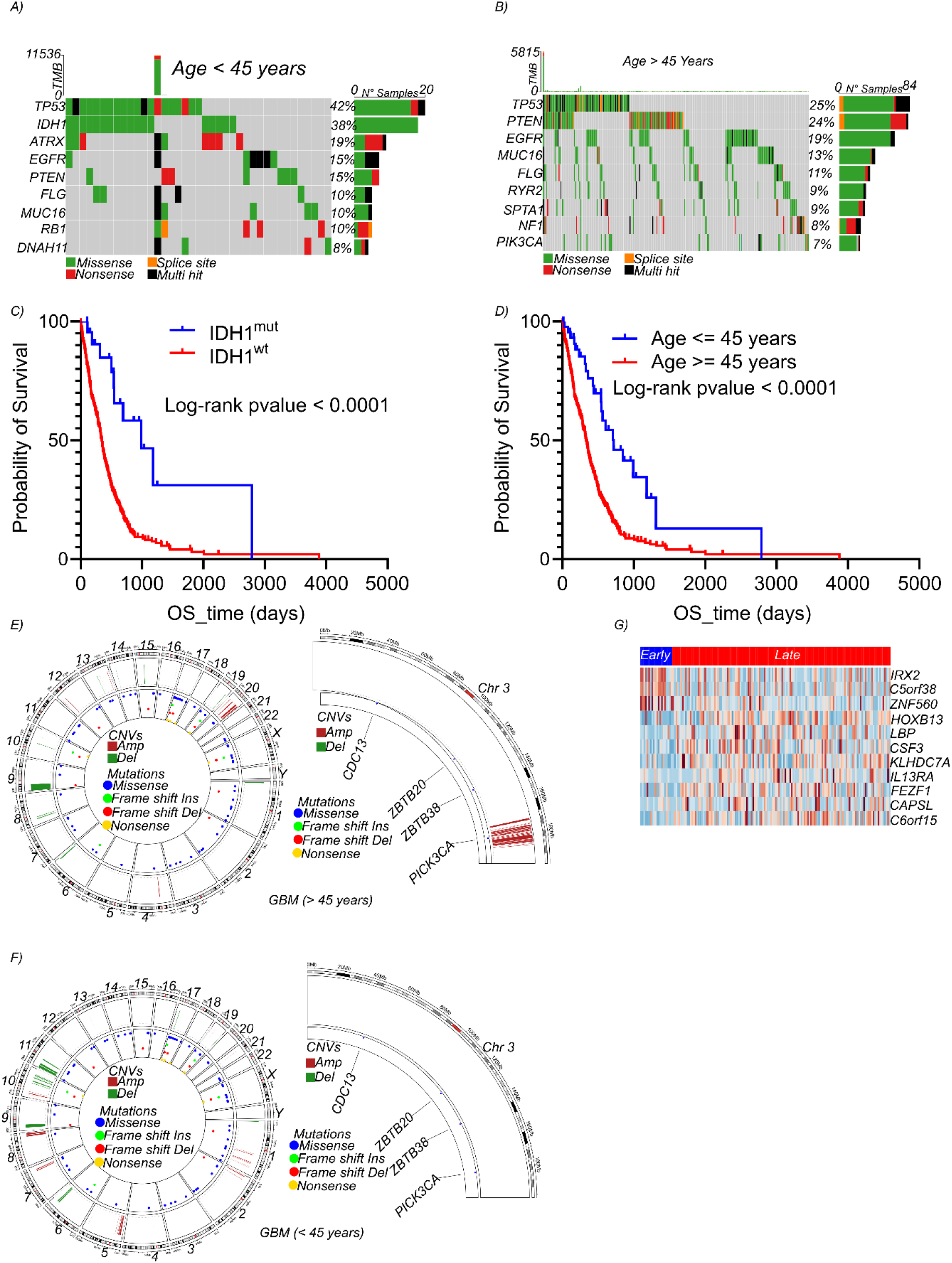
Early onset of glioblasoma is associated with IDH1 and TP53 mutations. **A**) An oncoprint showing the top 20 most mutated genes in glioblastoma in patients diagnosed with glioblastoma before the age of 45 years. **B**) An oncoprint showing the top 20 most mutated genes in glioblastoma in patients diagnosed with glioblastoma after the age of 45 years. **C**) A Kaplan-Meier overall survival curve for glioblastoma patients with and without *IDH1* mutation. **D**) A Kaplan-Meier overall survival curve for glioblastoma patients diagnosed before and after the age of 45 years. **E**) A circos plot showing somatic mutations and copy number alteration in glioblastoma patients diagnosed after the age of 45 years (left panel) and a zoom on chromosome 3 showing amplification at the *SOX2* gene locus (right panel). **F**) A circos plot showing somatic mutations and copy number alteration in glioblastoma patients diagnosed before the age of 45 years (left panel) and a zoom on chromosome 3 showing no amplification at the *SOX2* gene locus (right panel) as seen in older patients. **G**) A heat map showing genes that are differentially expressed between glioblastoma patients diagnosed before and after the age of 45 years.

### *SOX2* is amplified in aggressive tumors

We identified and annotated all common sCNA in the 10 most aggressive cancers. (**Supplementary table 1 & 2**). Differential gene expression analyses comparing the 10 most aggressive and seven less aggressive cancers was then performed (**supplementary table 3**). Both data sets were intersected to find gene dosage alterations affecting gene expression (**supplementary table 4**). There were more than 50 genes with somatic copy number amplification that were transcriptionally upregulated (**figure 3a**) and only less than 5 genes with sCNdel that were transcriptionally downregulated. To identify the major players involved in tumor aggressiveness, we classified all affected genes according to their importance in the aggressive disease phenotype using the Boruta package. As shown in **figure 3b**, *SOX2* was the most important variable, followed *by AADAC, FAM83A, UPK1B* and *MYEOV.* Multivariate cox proportional regression performed for the top 20 most important genes revealed, that high expression of the genes *SOX2, AADAC, FAM83A, MYEOV, UPKB1, ZIC5, ARL14, LYPD2, CALB1* and *MAGEA12* were associated with poor survival across all 27 cancer entities analyzed (**figure 4a**). Individually, Kaplan-Meier analysis of these genes revealed strong association with patient overall survival in all 27 cancer entities (**figure 4b**). We investigated if *SOX2* might indeed be regulating the expression of the identified genes. Using published *SOX2* ChIP-seq data (ENDCODE), we found *SOX2* peaks around three of these genes (*AADAC*, *FAM83A* and *MYEOV* (**figure 4c & 4D**), meanwhile majority of *SOX2* peaks were not promoter associated but either within introns or intergenic, suggesting possible enhancer binding (**figure 4e**). It is therefore very plausible, that *SOX2* regulates a gene network driving aggressive phenotypes.

**Figure 3:**
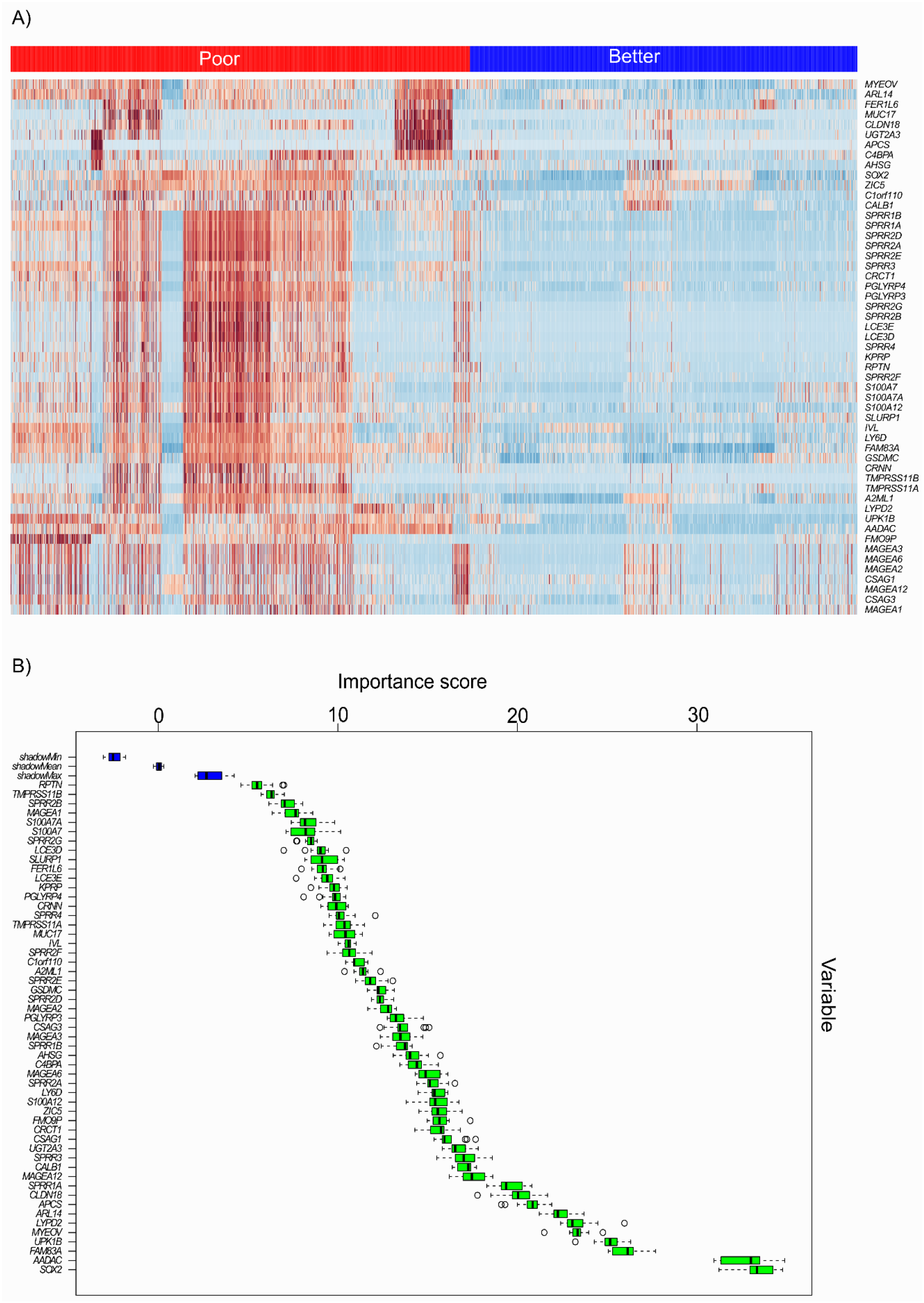
Copy number amplification drives concomitant transcriptional upregulation of genes modulating tumor aggressiveness. **A**) A heat map showing all genes with copy number amplifications and concomitant upregulation (log fold change ≤ 0.9, FDR < 0.05) in the top 10 most aggressive tumors. **B**) A variable of importance box plot for all gene with copy number amplifications and concomitant upregulation in the top 10 most aggressive tumors.

**Figure 4:**
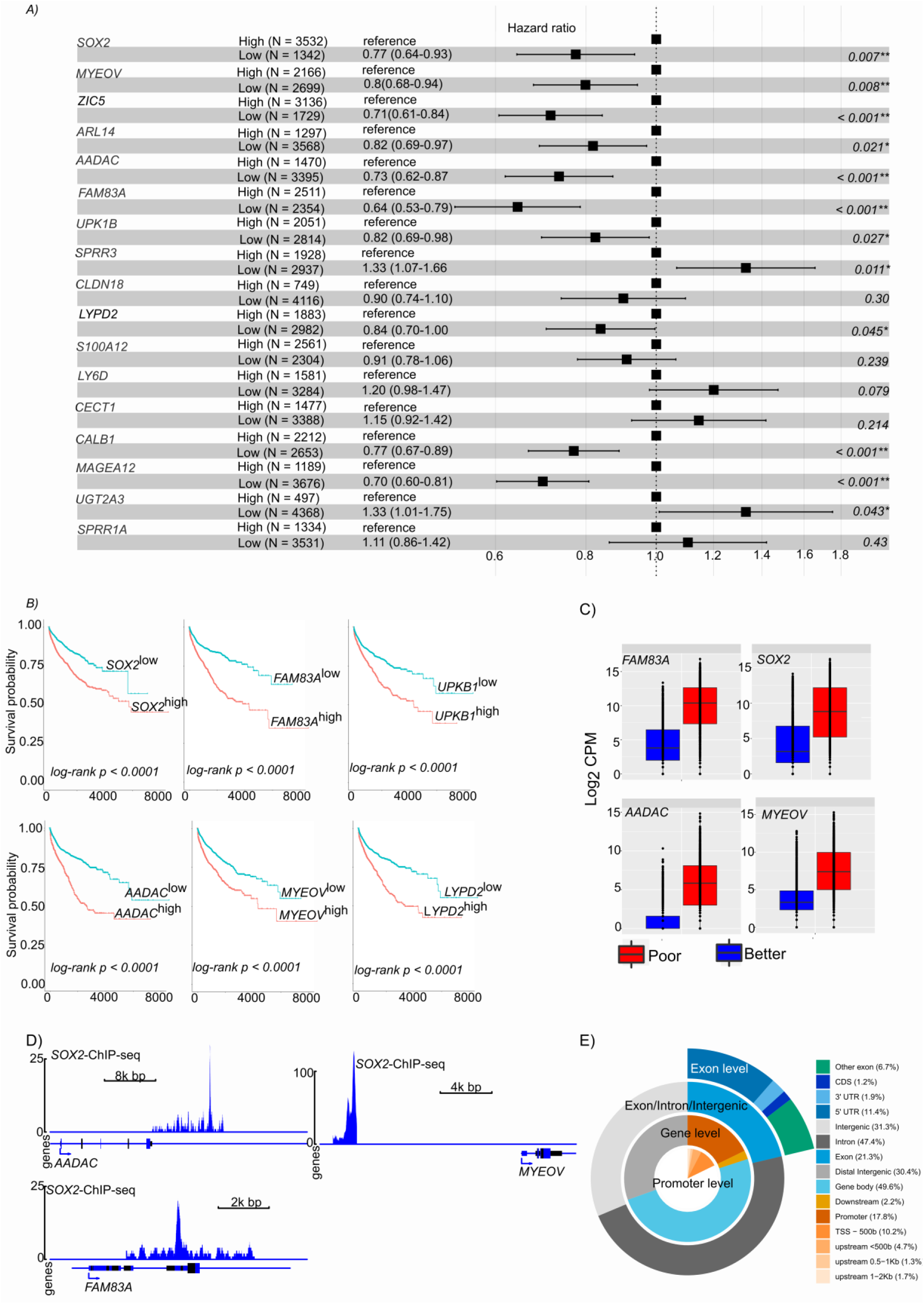
Genes with somatic copy number amplifications and concomitant upregulation are associated with patient outcome. **A**) A forest plot showing multivariate cox proportional hazards regression for genes with somatic amplification and transcriptional upregulation in poor outcome cancers. **B**) Kaplan-Meier overall survival plots for the top most significant genes in the multivariate model. **C**) Boxplots showing the transcript expression of SOX2-regulated genes. These genes are copy number amplified in poor outcome cancer and show concomitant upregulation. D) Accumulation of SOX2 peaks around its targets genes amplified and upregulated in poor outcome cancers. **E**) Feature plot, show the distribution of SOX2 ChIP-seq peaks across different genomic elements.

### *SOX2* enhances pro-inflammatory signaling via *FOSL2*

Gene set enrichment analysis of the hall mark of cancer revealed strong enrichment in pro-inflammatory pathways such as TNFA signaling via NFkB, *IL6-JAK-STAT3,* signaling, inflammatory response, interferon alpha/gamma response among others (**figure 5a & 5b**). Looking at the most enriched hallmark gene set (TNFA signaling via NFkB), the top 10 most enriched genes included two members of the *FOSL* transcription factor family (*FOSL1* and *FOSL2*, **figure 5c**). Given the implication of FOS genes in inflammation (32–34), we focused on these genes. Analysis of *SOX2* ChIP-seq data revealed that *SOX2* binds to the promoter of *FOSL2,* which can in turn regulate *IL6*, a key mediator of cellular inflammation (**figure 5d**). *SOX2* knockdown led to the downregulation of several genes, including *FOSL2* and several other HOX family transcription factors (**figure 5e, supplementary table 5**). To confirm the direct involvement of *SOX2* in oncogenic and inflammatory activities we performed pathway analysis on *SOX2* ChIP-seq data and observed a strong enrichment in pathways driving several cancer entities as well as NFkB-related inflammatory properties *(IL17* signaling pathway, **figure 5f**). Finally, we investigated if *FOSL2* is driving the enriched gene sets. To this end, we again performed gene set enrichment analysis in *FOSL2^high^* and *FOSL2^low^* (> 10000 and less than 1000 transcripts, respectively) tumor samples from all 27 cancer entities and compared the enriched hallmark gene sets. As shown in **figure 5g & supplementary table 6**, most of the gene sets enriched in the highly aggressive tumors were equally enriched in the *FOSL2^high^* tumors. There was equally a strong overlap between genes upregulated in aggressive tumors and in *FOSL2^high^* tumors (**figure 5h**).

**Figure 5:**
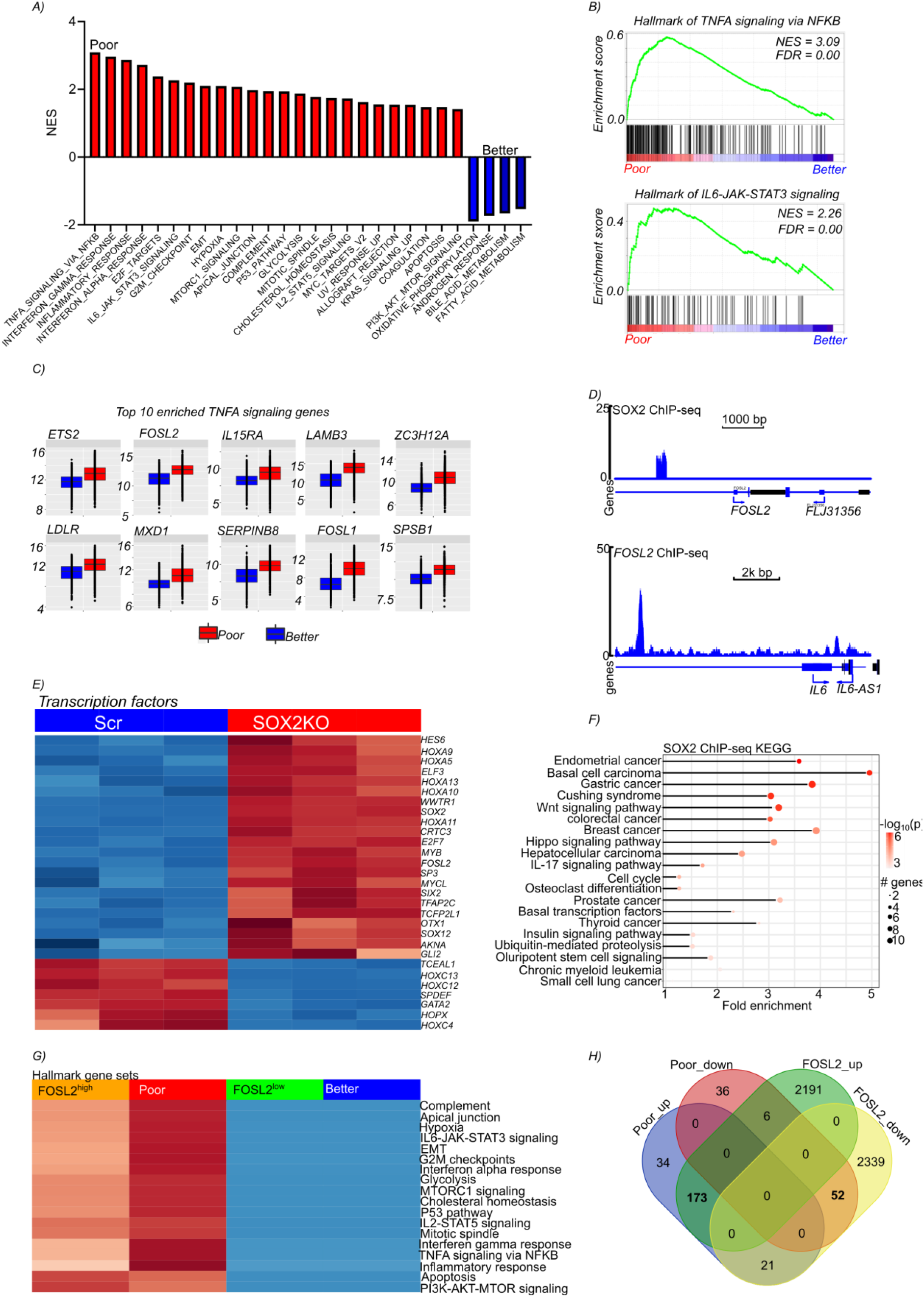
SOX2 promotes disease aggressiveness by enhancing inflammatory and oncogenic signaling via FOSL2. **A**) A bar plot showing enriched hallmark gene sets in poor and better outcome cancers. Gene sets are considered to be enriched, if the show a false discovery rate of < 0.05. **B**) Representative enrichment plot for pro-inflammatory hallmarks gene sets that are enriched in poor outcome cancers. **C**) Boxplots showing the 10 most enriched gene sets from the hallmarks of TNFA signaling via NFKB. The hallmarks of TNFA signaling via NFKB was the most significantly enriched hallmark gene set in poor outcome cancers. **D**) *SOX2* ChIP-sep peak profile showing enrichment around the *FOSL2* gene (upper panel) and *FOSL2* ChIP-seq peaks around the *IL6* gene (IL6-JAK-STAT3 signaling is one of the top enriched hallmark gene sets in poor outcome cancers). **E**) A heatmap showing the expression of significantly differentially expressed transcription factors upon SOX2 knockdown. **F**) A pathway plot showing significantly enriched pathways in *SOX2* ChIP-seq data. The intensity of the color shows the strength of the enrichment p-value, while the size of the circle indicate the number of enriched genes. **G**) A heatmap comparing enriched hallmark pathways in poor and better outcome cancers and *FOSL2*^high^ and *FOSL2^low^* samples. *FOSL2*^high^ and *FOSL2^low^* samples are samples with less than 1000 copies or more than 10000 copies, respectively. This approach was used investigate if high expression of FOSL2 is related to the enriched hallmark gene sets seen in the poor outcome cancers. H) A Venn diagram showing the intersection between upregulated and downregulated genes in poor out come and better outcome samples and between *FOSL2^hgh^* and *FOSL2^low^* samples.

### Inflammation induces immunosuppression and Cytidine deamination

We observed a high expression of *IL6* and its downstream effector *STAT3* in aggressive tumors and in *FOSL2^high^* tumors (**figure 6a**). We used gene expression data from laser microdissected PDAC samples to determine the tumor compartment driving the observed inflammation. High expression of *FOSL2* and *STAT3* were observed in the stroma (**Figure 6b**). *IL6* expression was very low in the data set and is not presented. We used PDAC, because of its aggressiveness and presence of dense fibrotic stroma. We then analyzed the expression of *IL6* in grade II and grade III PDAC (**figure 6c, left panel**) as well as in quasi-mesenchymal and classical PDAC (**figure 6c, right panel**). As expected, *IL6* was significantly upregulated in grade III as well as in quasi-mesenchymal PDAC tumors. We then investigated the expression of immune cell markers in our cohort of almost 300 samples including 41 normal pancreas tissue, 59 chronic pancreatitis tissue and 195 resected PDAC tissues. We observed a steady increase in the expression of *FOXP3, LRRC32,* as well as *FOSL2* and *IL6* from normal through pancreatitis to PDAC (**figure 6d**). Given that inflammation is associated with immunosuppression, we evaluated the immune cell proportion in the different PDAC samples and observed significant increase in Tregs, proinflammatory M1 macrophages and neutrophils, while activated NK cells were decreased from normal pancreas tissue through pancreatitis to PDAC (**figure 6e**). It is known, that prolonged inflammation induces cytosine deamination as well as DNA double strand breaks and leads to genome instability. We therefore analyzed mutational signatures in aggressive and less aggressive tumors. We observed an enrichment in cytosine deamination mutational signatures in the aggressive tumors (**figure 6f**), with a characteristic C>G mutations associated with inflammation. Mutational analysis revealed high rates of *TP53* mutations in the aggressive tumors (**figure 6g**). As in the PDAC data, immune cell infiltration analysis of aggressive and less aggressive tumors revealed similar infiltration patterns (**figure 6h**).

**Figure 6:**
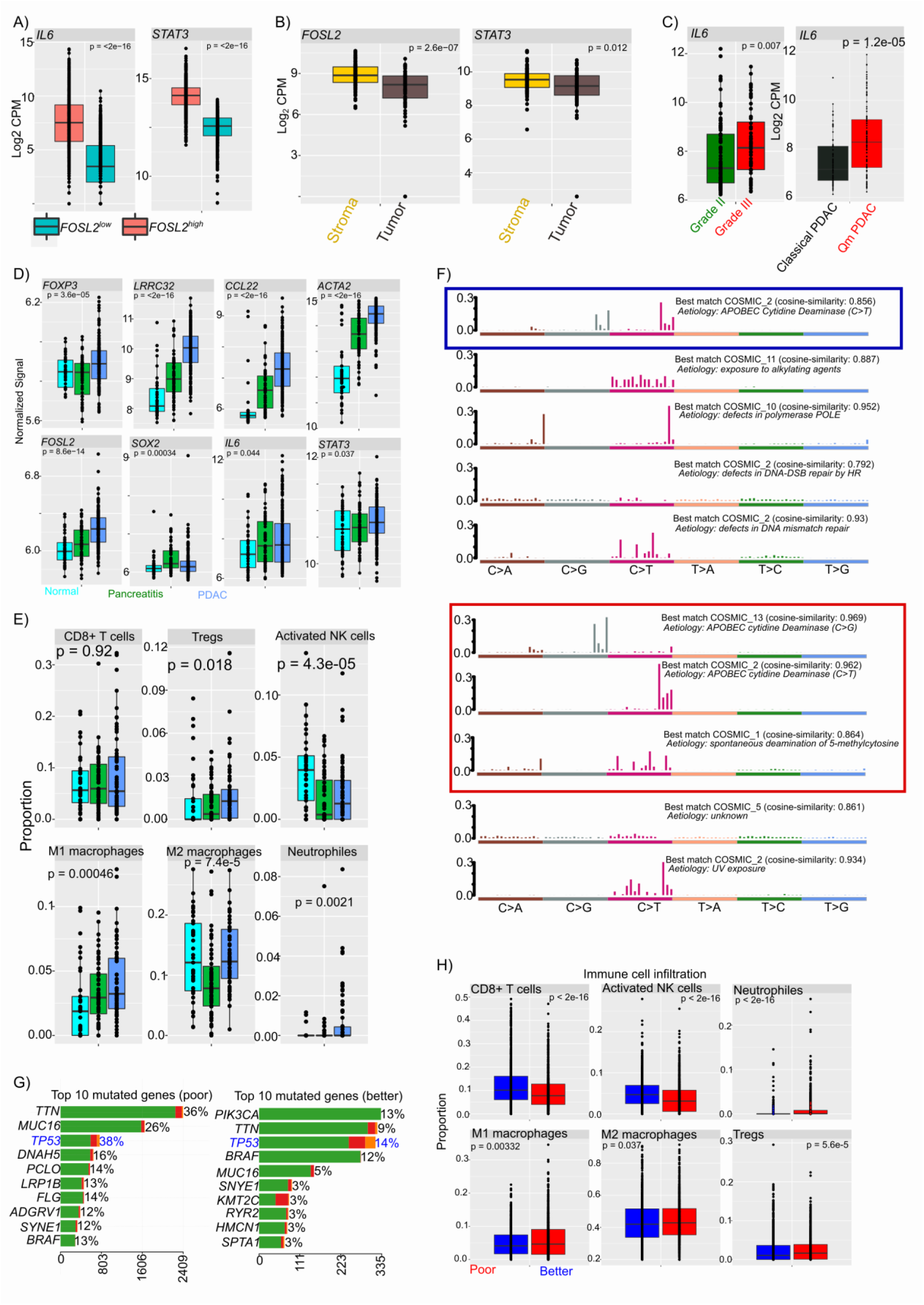
Proinflammatory tumor microenvironment induces characteristic cytidine deamination mutational signatures in aggressive tumors. **A**) Boxplots showing the expression of the proinflammatory cytokine *IL6* and its downstream effector *STAT3* in *FOSL2* high and low tumors. *FOSL2* high and low groups are the same as described above. **B**) Expression of FOSL2 and the proinflammatory mediator STAT3 in different compartments of PDAC tumors. Data is derived from laser microdissected (Maurer *et al*., 2019) PDAC tumors. The expression of *IL6* was very low in majority of the cases and is not presented. **C**) IL6 gene expression in grade II and grade III PDAC tumors (left panel) and in Qm and classical PDAC subtypes (right panel). Data was generated from resected PDAC samples. **D**) Expression of inflammatory mediators and tumor microenvironment marker genes in normal pancreas (n = 41), chronic pancreatitis tissue (n = 59) and PDAC tissues n = 195). **E**) Immune cell proportions derived from deconvolution of gene expression data from normal pancreas tissue, chronic pancreatitis tissue and PDAC tissue. Immune cell proportions were determined with CIBERSORT as implemented in TIMER. **F**) Top five mutational signatures in cancer entities with better outcome (upper panel) and poor outcome (lower panel). **G**) Top ten most mutated genes in poor outcome cancers 8left panel) and better outcome cancers (right panel). **H**) Boxplots showing immune cell proportions in poor outcome vs better outcome cancers,

## Discussion

We investigated the molecular drivers of cancer aggressiveness across 27 cancer using data from the TCGA and other publicly available data sets. We observed a significant association between sCNA and cancer fatality rate. In effect, somatic copy number alterations fuels genetic heterogeneity and supports the amplification of oncogenes and drug resistance genes in cancer (35, 36), thereby influencing outcome. Tumor mutational burden was not associated with tumor lethality, except in SKCM, as it has was previously reported in another pan-cancer analysis (37). We observed more than 15% of GBM in cases aged below 45 years. There was however, no significant difference in sCNA and TMB in GBM patients of the different age groups. Somatic mutations analyses revealed high levels of *TP53*, *IDH1* and *ATRX* mutations in younger patients, while late onset was characterized amplification of dedicated oncogenes such as *PIK3CA.* In effect, *IDH1* mutation in GBM have been reported to be associated with better prognosis (30, 31). Given the association of *IDH1* mutations with younger patients, it is very likely, that age contribute to the observed survival benefit. This observation therefore plaid in favor of age consideration in subsequent translational studies and clinical trials, especially for targeted therapies.

Integrated multi-omic analysis identified the transcription factor, *SOX2* to be amplified and transcriptionally upregulated in aggressive cancers. *SOX2* has previously been implicated with cancer development (38), stemness and pluripotency (39, 40). Apart from *SOX2*, other genes involved in cancer aggressiveness such as *ZIC5*, *FAM83A*, *AADAC* and *MYEOV* were among the top most significantly associated genes and were equally associated with patient outcome. The non-coding RNA transcript *MYEOV* has been shown to drive disease aggressiveness in cancer (41, 42). Furthermore, *FAM83A* has been associated with enhanced tumor cell proliferation and metastasis by regulating Wnt/ß catenin signaling (43) and promotes tumorigenicity via the ERK, PIK3/AKT/mTOR pathway (44). Interestingly, we observed *SOX2* binding events around some of these genes, suggestive of a *SOX2*-regulated gene network.

Pathway analyses revealed activation of several proinflammatory pathways such as *TNFA* signaling via NFkB, inflammatory response and *IL6-JAK-STAT3* signaling among other in aggressive tumors. Two gene of the *FOSL* transcription factor family, which was recently implicated in inflammation (34), were among the top enriched genes in the TNFA signature. In effect, TNFA signaling is a prominent proinflammatory signaling pathway (45). Investigating possible associations between *SOX2* and the *FOSL* genes, we observed *SOX2* regulatory activity on *FOSL2,* and the later could in turn regulate *IL6* gene expression, indicating that *IL6* is possibly downstream of *FOLS2,* which itself is a target of *SOX2.* Pathway analysis on SOX2 ChIP-seq data revealed activation of pathways driving the development of several cancers as well as pro-inflammatory pathways. These observations suggest the implication of *SOX2* in sustaining inflammatory processes in aggressive tumors via *FOSL2* and *IL6*. Tumors with high *FOSL2* expression shared similar pathway activation as aggressive tumors, strengthening the interconnection between *FOSL2* and cancer aggressiveness.

Aggressive tumors equally showed overexpression of proinflammatory cytokines and higher levels of immunosuppression. Prolonged inflammation, principally driven by NFkB can activate cytidine deamination (28) and lead to a characteristic mutational signature and DNA damage. Indeed, we observed a strong enrichment of cytidine deamination mutational signatures in aggressive tumors, a direct indicator of prolonged inflammation in these tumors. Additionally, *TP53* mutations were more predominant in aggressive tumors. It is plausible, that inflammation-induced DNA damage affect tumor suppressor gene function, thereby promoting genomic instability and releasing the cell cycle brake to unleash uncontrolled proliferation.

## Conclusion

Taken together, our data uncover the implication of a *SOX2*-regulated gene expression network controlling cancer aggressiveness via *FOSL2* by activating and sustaining pro-inflammatory TME leading to DNA damage and genomic instability. Targeting these pro-inflammatory processes might therefore indeed minimize DNA damage and improve patient outcome.

## Declarations

### Ethics approval and consent to participate

Not applicable

### Consent for publication

Not applicable

### Availability of data and materials

All data used in this manuscript is publicly available through the TCGA or under the given accession number from the GEO. Data supporting our findings are reported in the manuscript and supplementary information

### Competing interests

The authors declare that they have no competing interests

### Funding

Not applicable

### Authors’ contributions

Study design, data acquisition, data analysis and manuscript draft (AJN and SSL), data curation AAZT, RK, EHMDB, SNA, manuscript editing and project resources GS, JH and JS

## Acknowledgements

Not applicable

## List of abbreviations

ACC: Adrenocortical carcinoma
BLCA: Bladder Urothelial Carcinoma
LGG: Brain Lower Grade Glioma
BRCA: Breast invasive carcinoma
CESC: Cervical squamous cell carcinoma and endocervical adenocarcinoma
CHOL: Cholangiocarcinoma
COAD: Colon adenocarcinoma
ESCA: Esophageal carcinoma
GBM: Glioblastoma multiforme
HNSC: Head and Neck squamous cell carcinoma
KICH: Kidney Chromophobe
KIRC: Kidney renal clear cell carcinoma
KIRP: Kidney renal papillary cell carcinoma
LIHC: Liver hepatocellular carcinoma
LUAD: Lung adenocarcinoma
LUSC: Lung squamous cell carcinoma
OV: Ovarian serous cystadenocarcinoma
PAAD: Pancreatic adenocarcinoma
PRAD: Prostate adenocarcinoma
SARC: Sarcoma
SKCM: Skin Cutaneous Melanoma
STAD: Stomach adenocarcinoma
TGCT: Testicular Germ Cell Tumors
THYM: Thymoma
THCA: Thyroid carcinoma
UCEC: Uterine Corpus Endometrial Carcinoma
UVM: Uveal Melanoma
EMT: epithelial to mesenchymal transition
TNFA: tumor necrosis factor alpha
IL: interleukin
TMB: tumor mutational burden
sCNA: somatic copy number alteration
sCNAmp: somatic copy number amplification
sCNdel: somatic copy number deletion
TCGA: the cancer genome atlas
GEO: gene expression omnibus
FDR: false discovery rate

---

**Figure S1:**
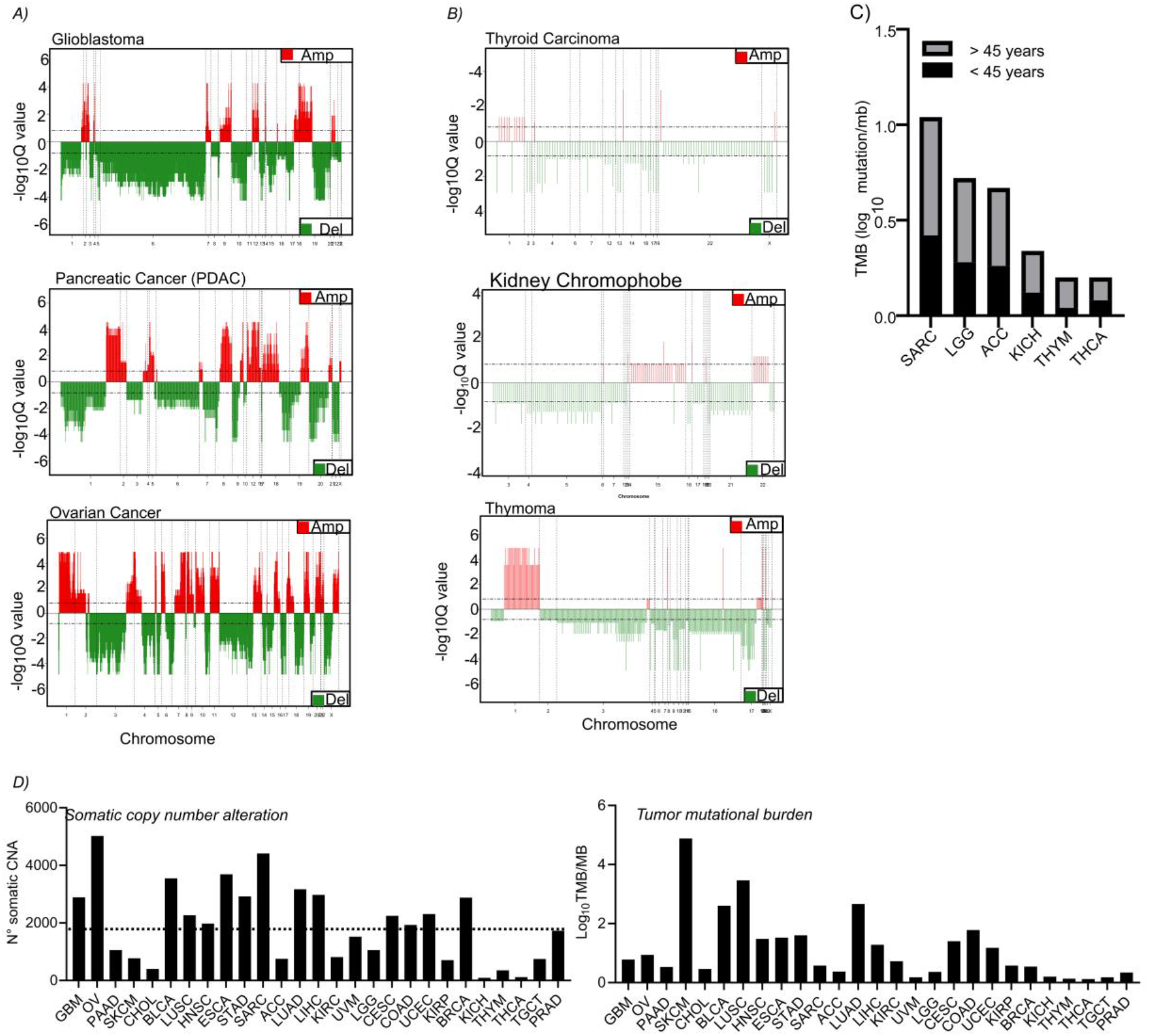
Highly aggressive cancers show higher numbers of CNAs and slightly higher tumor mutational burden. **A**) CNA plots for top three most aggressive and prevalent cancer. **B**) CNA plots for top three least aggressive and prevalent cancers. **C**) Bar plot showing tumor mutational burden for different age groups for all cancer with more than 10% of cases diagnosed before 45 years of age except those shown in previous figures. **D**) A bar plot show the number of CNAs identified in each of the analyzed cancer entities (left panel) and a bar plot show the tumor mutational burden in each of the analyzed cancer entities (right panel).

**Figure S2.**
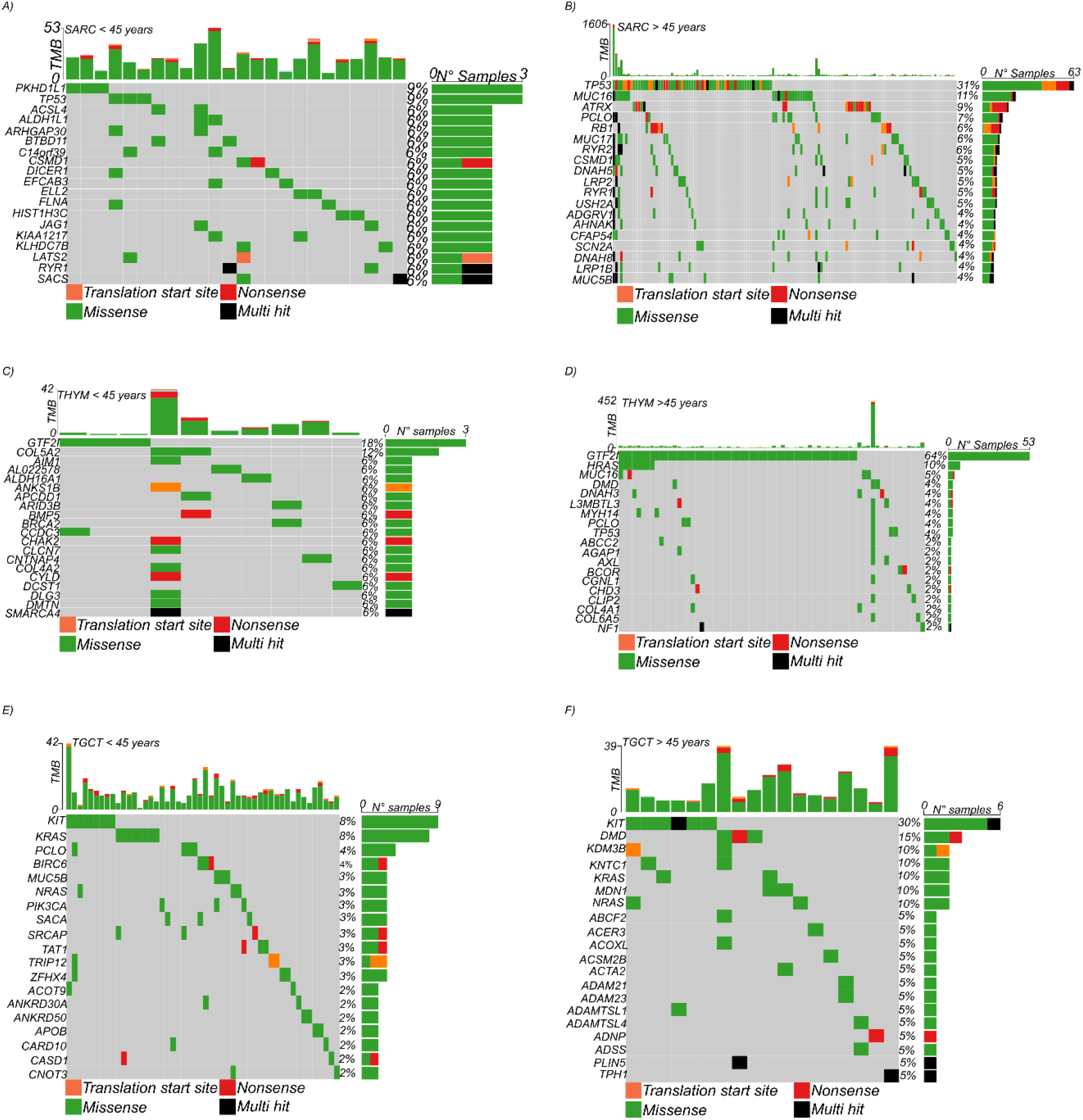
Somatic gene mutations are prevalent in elderly patients than younger patients. **A**) An oncoprint showing somatic mutations in sarcoma patients diagnosed before the age of 45 years. **B**) An oncoprint showing somatic mutations in sarcoma patients diagnosed after the age of 45 years. **C**) An oncoprint showing somatic mutations in thymoma patients diagnosed before the age of 45 years. **D**) An oncoprint showing somatic mutations in thymoma patients diagnosed after the age of 45 years. **E**) An oncoprint showing somatic mutations in patients with testicular germ cell tumors diagnosed before the age of 45 years. **F**) An oncoprint showing somatic mutations in patients with testicular germ cell tumors diagnosed after the age of 45 years.

## Notes

### Competing Interest Statement

The authors have declared no competing interest.

